# Rev-RRE activity modulates HIV-1 replication and latency reactivation: Implications for viral persistence and cure strategies

**DOI:** 10.1101/2025.01.06.631466

**Authors:** Godfrey A. Dzhivhuho, Patrick E. H. Jackson, Ethan S. Honeycutt, Flavio da Silva Mesquita, Jing Huang, Marie-Louise Hammarskjold, David Rekosh

## Abstract

The HIV-1 Rev-RRE regulatory axis plays a crucial role in viral replication by facilitating the nucleo-cytoplasmic export and expression of viral mRNAs with retained introns. In this study, we investigated the impact of variation in Rev-RRE functional activity on HIV-1 replication kinetics and reactivation from latency. Using a novel HIV-1 clone with an interchangeable Rev cassette, we engineered viruses with different Rev functional activities and demonstrated that higher Rev-RRE activity confers greater viral replication capacity while maintaining a constant level of Nef expression. In addition, a low Rev activity virus rapidly acquired a compensatory mutation in the RRE that significantly increased Rev-RRE activity and replication. In a latency model, proviruses with differing Rev-RRE activity levels varied in the efficiency of viral reactivation, affecting both initial viral release and subsequent replication kinetics. These results demonstrate that activity differences in the Rev-RRE axis among different viral isolates have important implications for HIV replication dynamics and persistence. Importantly, our findings indicate that bolstering Rev/RRE activity could be explored as part of latency reversal strategies in HIV cure efforts.

**Author Summary:** The activity of the HIV-1 Rev-RRE axis is essential for viral replication and varies among primary viral isolates. However, the role of this for viral fitness, evolution, and persistence has not previously been investigated. Our results show that during *in vitro* replication, there is a selective fitness advantage for viruses with higher Rev-RRE activity and that HIV has the ability to fine tune this regulatory system with minimal sequence changes. Additionally, the maintenance of Nef expression in low Rev activity viruses suggests a potential mechanism for balancing immune evasion and replication capacity in different selection landscapes within a host. We also show that viruses with low Rev-RRE activity are more difficult to reverse from latency than viruses with higher Rev-RRE activity. Thus, differences in provirus Rev-RRE activity may be a barrier to developing effective latency reversal strategies. These findings provide new insights into the complex roles that the Rev/RRE axis plays in functionality, viral fitness, evolution, and persistence.

## Introduction

Retroviruses have compact genomes and utilize a complex alternative splicing pattern to express the full range of viral transcripts required for replication. As a result of this strategy, viral RNAs with retained introns (IR-mRNAs) have to be exported from the cell nucleus to the cytoplasm for genome packaging into new virus particles and for production of structural proteins. Retroviruses use one of two strategies to overcome the normal cellular restrictions to export of these RNAs. “Simple” retroviruses, such as the Mason-Pfizer Monkey Virus, contain an RNA structure in the IR-mRNAs, the CTE (1), that directly recruits the cellular factors Nxf1 and Nxt1 to permit nuclear export. In contrast, “complex” retroviruses require both a structured element in the IR-mRNAs and a viral regulatory protein, to accomplish this task (2).

HIV-1 is a complex retrovirus that utilizes the viral protein Rev, and an RNA structure known as the Rev Response Element (RRE), to export its viral IR-mRNAs (3–5). Rev is translated from a completely spliced mRNA and is imported into the nucleus, where it binds to the RRE present in the unspliced and incompletely spliced viral mRNAs. Rev then acts as an adapter on the RRE to recruit cellular factors including Crm1 and RanGTP. This creates a complex that enables the export of the IR-mRNA to the cytoplasm (6–11).

The RRE of the lab isolate pNL4-3 that used in many *in vitro* studies, is a highly structured element of about 350 nucleotides (12). The element exists in a dynamic equilibrium between two low-energy conformations containing either four or five stem loops (13). The highest affinity binding site for Rev is located on stem-loop IIB, and a complex of about six to eight Rev monomers sequentially assembles on the RRE (14). As with all other regions of the HIV genome, the RRE is subject to frequent mutations, resulting in sequence differences among primary isolates. We previously observed the sequence, structural, and functional evolution of the RRE in circulating viral variants in individuals over time (15, 16). A small number of nucleotide changes was sufficient to cause a substantial alteration in RRE secondary structure, as well as significant differences in Rev-RRE functional activity (16). Previous studies have also shown that one or two changes in the RRE allows the virus to escape inhibition by small molecules or a trans-dominant negative Rev protein (17, 18).

The Rev protein is also subject to significant sequence and functional variation. The prototypical (subtype B) Rev consists of 116 amino acids and includes a bipartite oligomerization domain (19), a nuclear export signal (20), and an arginine-rich motif, which serves as both the RNA binding region and the nuclear localization signal (21, 22). We previously described two subtype G Revs from primary isolates that showed widely disparate activity levels (23). The high activity, 9-G, and low activity, 8-G, Rev sequences differed at 29 amino acid positions. However, the activity difference between the two Revs was subsequently shown to be attributable to two single amino acid differences in the first oligomerization domain (24).

HIV replication is dependent on a functional Rev-RRE axis, but the functional plasticity of Rev and the RRE permits fine-tuning of Rev-RRE activity through small sequence changes in either element. The consequences of this variation for HIV pathogenesis have not been fully explored. However, in another complex retrovirus, Equine Infectious Anemia Virus (EIAV), variations in the Rev-RRE axis over time are clearly associated with clinical disease progression (25–27). In the case of HIV, Rev activity has been shown to be a determinant of the sensitivity of HIV-1-infected primary T cells to cytotoxic T-lymphocyte killing (28) and RRE activity has been positively correlated with the rate of CD4 count decline (29, 30). Rev-RRE activity has also been shown to affect the fitness of viral variants in the context of female-to-male sexual transmission (31). Thus, it is reasonable to hypothesize that Rev-RRE activity differences may play a role in many aspects of HIV pathogenesis, including in the dynamics of the reservoir and viral rebound after treatment interruption. In this study, we explored the effect of Rev-RRE activity differences on viral replication kinetics and *in vitro* fitness in the context of both a spreading infection and reactivation from latency.

## Results

### Construction and characterization of an HIV proviral clone with an interchangeable Rev

The coding region of the HIV Rev gene overlaps the coding regions of both Env and Tat. Thus, to investigate the impact of variation in Rev activity on viral replication kinetics, it was essential to create an infectious clone that relocated Rev into a position where it could be manipulated without changing other HIV genes. To do this, we first silenced Rev in the pNL4-3 proviral clone by mutating the AUG initiation codon to ACG and the 23^rd^ codon, UAU, to a stop codon, UAA. The stop codon in Rev introduced a conservative serine to threonine change at amino acid position 70 in the overlapping Tat open reading frame (Fig 1A). We then inserted a cassette upstream of Nef that contained a cDNA copy of NL4-3 Rev, followed by an internal ribosomal entry site (IRES). This allowed both Rev and Nef to be expressed from a fully spliced HIV mRNA that would normally encode only Nef.

**Fig 1.**
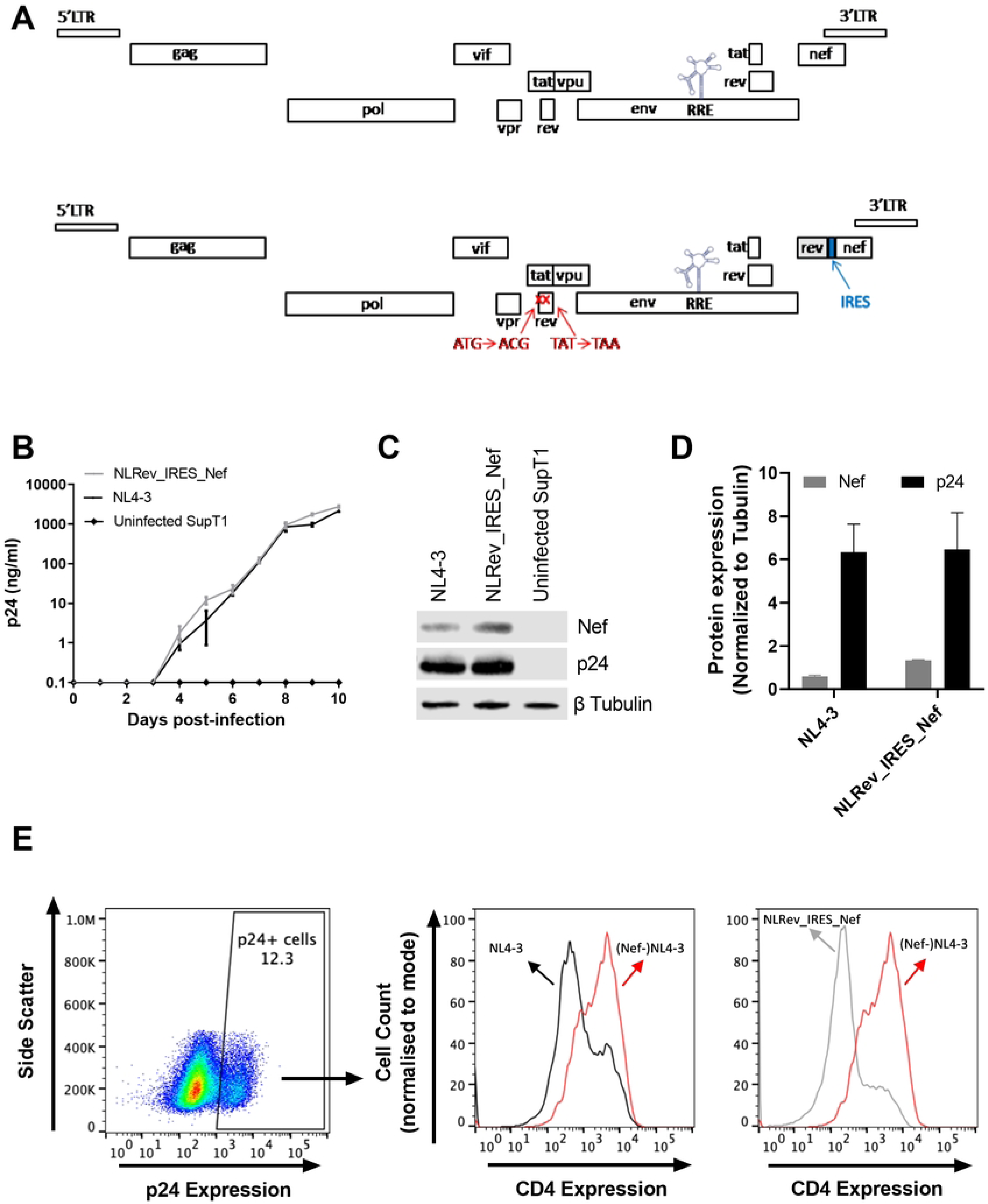
Construction and characterization of an HIV proviral clone with an interchangeable *rev* gene. (A) Schematic of HIV proviral constructs. Upper panel: original NL4-3. Lower panel: modified NLRev_IRES_Nef. The native *rev* gene was silenced by start codon mutation (AUG to ACG) and stop codon introduction at position 23 (UAC to UAA). An NL4-3 *rev* cDNA cassette with an IRES was inserted upstream of *nef*, flanked by restriction sites for easy exchange. (B) SupT1 cells were infected with NL4-3 (black line) and NLRev_IRES_Nef (grey line) viruses at an MOI of 0.005. Viral replication was monitored by p24 ELISA of culture supernatants over time. (C) Representative Western blot analysis of p24 and Nef expression in SupT1 cells infected with NL4-3 or NLRev_IRES_Nef viruses at day 8 post-infection. Beta tubulin was used as a loading control. (D) Quantification of Nef (grey bar) and p24 (black bar) expression from Western blot analysis. (E) SupT1 cells were infected with NL4-3 (black), NLRev_IRES_Nef (grey), and Nef-negative NL4-3 control (red) viruses at an MOI of 0.005 for 8 days. Left panel: Representative flow cytometry plots of intracellular p24 staining in infected cells. Right panels: Quantification of surface CD4 expression in infected cells. Error bars represent mean ± SEM of three independent experiments for panel B, and mean ± SD of three independent experiments for panels D.

Another important feature of this construct is that the Rev sequence is flanked by restriction enzyme sites so that it can easily be exchanged for alternative Rev sequences. This allowed us to make two additional constructs using sequences from two Rev variants (8-G, accession FJ389367; 9-G, accession JX140676). The 8-G and 9-G Rev proteins were previously shown to have disparate functional activities. When tested on the NL4-3 RRE, the functional activity of these Revs differed by 4-5-fold (23). The new proviral clones were named pNLRev_IRES_Nef (HamRek archive #5833), p8-GRev_IRES_Nef (HamRek archive #5830) and p9-GRev_IRES_Nef (HamRek archive #5831).

To analyze how moving Rev to a different region affected replication and Nef expression, we first transfected 293T cells with pNLRev_IRES_Nef and the original pNL4-3 proviral clone. Forty-eight hours (48h) after transfection, supernatants were collected and assayed for p24 (S1 Fig1B). One hundred nanograms of supernatant p24 were then used to infect SupT1 cells (1x10^6^/ml), creating intermediate viral stocks (S1 Fig 1C). These viral stocks were passaged a second time in SupT1 cells (S1 Fig !D), creating the final expanded viral stocks whose titers were determined by TCID_50_. The details of this procedure are shown in S1 Fig 1A and further described in materials and methods.

We first compared the replication kinetics of the two viruses. To do this, parallel cultures of SupT1 cells were infected with both viruses at a multiplicity of infection (MOI) of 0.005. Viral replication was assessed by measuring the levels of p24 in culture supernatants over time. As can be seen in Fig 1B, these viruses replicated similarly as measured by p24 production.

To analyze Nef expression, cells were collected eight days post-infection and subjected to Western blot analysis. Whereas the levels of p24 were comparable between the two viruses, the NLRev_IRES_Nef virus appeared to express a slightly higher level of Nef compared to NL4-3 (Fig 1C and 1D). This could be due to Nef expression from both spliced and unspliced mRNA in the engineered construct because of the presence of an IRES. We next assessed Nef functionality in these two viruses. HIV Nef downregulates cell surface CD4 expression through clathrin- and adaptor protein complex 2-dependent endocytosis (32, 33). SupT1 cells were infected with NL4-3 or NLRev_IRES_Nef viruses at an MOI of 0.005 and then stained for intracellular p24 and surface CD4 expression at 8 days post-infection. Cells infected with a Nef-negative NL4-3 virus were used as a control. Both NL4-3 and NLRev_IRES_Nef-infected cells displayed downmodulation of CD4 compared to the Nef-negative control, demonstrating that the modified viruses express a functional Nef protein (Fig 1E).

Taken together the results of these experiments indicate that the relocation of Rev and the incidental change in Tat did not impair viral replication kinetics or the ability of the virus to produce a functional Nef protein.

### Viruses with low Rev functional activity replicate poorly compared to viruses with high Rev functional activity

We next examined the replication kinetics of the 8-GRev_IRES_Nef and 9-GRev_IRES_Nef viruses in comparison with NLRev_IRES_Nef. To do this, SupT1 cells were infected with each virus from the expanded stocks at an equal MOI (0.005). Measurement of p24 in culture supernatants by ELISA revealed that the 9-GRev_IRES_Nef virus replicated similarly to the NLRev_IRES_Nef virus. However, the 8-GRev_IRES_Nef virus replicated much less efficiently (Fig 2A).

**Fig 2.**
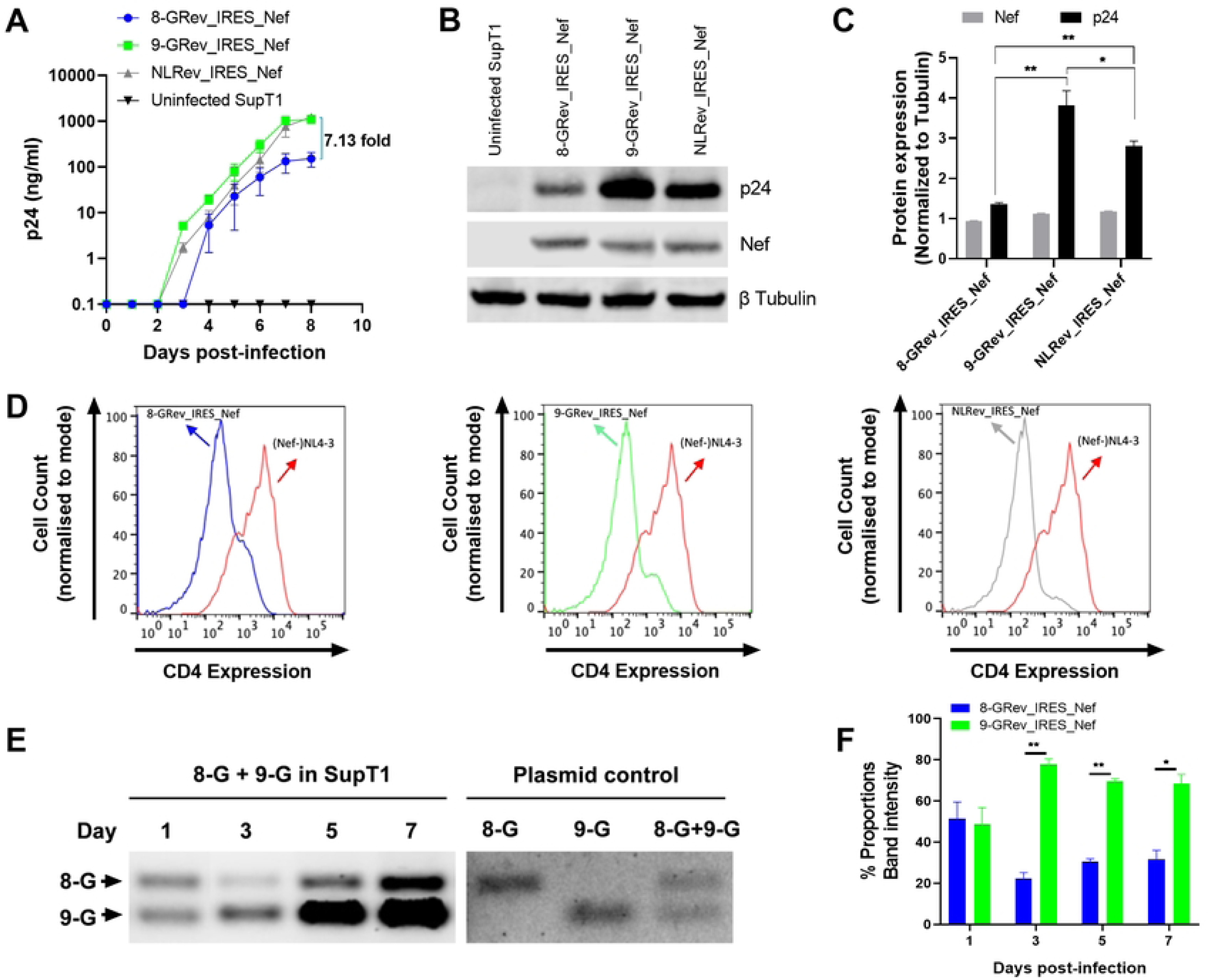
Viruses with low Rev functional activity replicate poorly compared to viruses with high Rev functional activity. (A) SupT1 cells were infected with 8-GRev_IRES_Nef (low Rev activity) (blue), 9-GRev_IRES_Nef (high Rev activity) (green), and NLRev_IRES_Nef (control) (grey) viruses at an MOI of 0.005. Viral replication was assessed by p24 ELISA of culture supernatants over time. (B) Western blot analysis of Nef and p24 expression in infected cells at day 8 post-infection. Beta-tubulin was used as a loading control. (C) Quantification of Nef and p24 expression from Western blot analysis in B, normalized to beta-tubulin. (D) Quantification of CD4 downmodulation in SupT1 cells infected with 8-GRev_IRES_Nef, 9-GRev_IRES_Nef, NLRev_IRES_Nef viruses at an MOI of 0.005 for 8 days, relative to Nef-negative NL4-3 virus. (E) SupT1 cells were co-infected with 8-GRev_IRES_Nef and 9-GRev_IRES_Nef viruses at an MOI of 0.005 each. Left panel: Representative gel image showing PCR products of cellular DNA targeting Rev at days 1, 3, 5, and 7 post-infection. Right panel: PCR plasmid controls. (F) Quantification of relative proviral DNA levels from the competition assay. Bars represent the ratio of 9-GRev_IRES_Nef to 8-GRev_IRES_Nef proviral DNA at each time point. Error bars represent ± SEM of three independent experiments for panel A, and ± SD of three independent experiments for panels C and F. Statistical analysis was performed using unpaired, two-tailed t-test with FDR correction for multiple comparisons (*p<0.05, **p<0.01).

Levels of Nef and p24 in the infected cells were also measured by western blot analysis. Although there was no difference in Nef expression, p24 levels were much lower in cells infected with the 8-GRev_IRES_Nef virus compared to those infected with NL4Rev-IRES-Nef or 9-GRev_IRES_Nef viruses. The 9-GRev_IRES_Nef virus exhibited the highest p24 expression (Fig 2B and 2C).

To determine whether Nef function was affected, the ability of each virus to downmodulate CD4 was tested. Flow cytometry of p24-stained cells revealed that all viruses, including the 8-GRev_IRES_Nef virus, downmodulated CD4 compared to the Nef-negative NL4-3 control (Fig 2D), showing that all the viruses expressed functional Nef.

To directly compare the replicative fitness of viruses with low (8-GRev_IRES_Nef) and high (9-GRev_IRES_Nef) Rev activity, a competition assay was performed by co-infecting SupT1 cells with both viruses. An MOI of 0.005 was used to ensure that the number of co-infected cells would be minimal and thus allow the activity of each Rev to be assessed in parallel. Cellular DNA was collected, and PCR was performed using primers targeting Rev, since the PCR products from the 8-GRev_IRES_Nef and 9-GRev_IRES_Nef viruses are distinguishable by size.

On day 1 both 8-GRev_IRES_Nef and 9-GRev_IRES_Nef proviruses were present in equal amounts, confirming the equal MOI used for infection (Fig 2E and 2F). On days 3, 5, and 7, a significantly higher number of proviral copies of the 9-GRev_IRES_Nef provirus were present in the co-infected cells, demonstrating that the high Rev activity virus outcompeted the low Rev activity virus over time.

### Continued passage of the low Rev activity 8-GRev_IRES_Nef virus selected for a single nucleotide change in the RRE

When we continued the infections shown in Fig 2A beyond day 8, we observed a dramatic increase in p24 production in cells infected with the 8-GRev_IRES_Nef virus (Fig 3A). We reasoned that this could be due to the selection of a viral mutant that replicated more efficiently. To examine this, both the RRE and Rev regions of integrated proviruses at the day 10 timepoint were amplified by nested PCR and sequenced using Sanger sequencing. No changes were observed in the Rev sequence. However, sequencing of the RRE from 8-GRev_IRES_Nef infected cells revealed that a considerable fraction of the proviruses displayed a single nucleotide change in the stem IIB region of the RRE (Fig 3B) at pNL4-3 residue 7214 (NL4-3 accession number U26942). Sequencing of proviral DNA from the day 6 cells also revealed the same change. In contrast, only negligible amounts of the C7214T mutation were found in proviral DNA from the day 1 cells. In addition, the mutation was completely missing in the proviruses present in the originally infected day 8 cells, the supernatant of which contributed the viral stock used for the subsequent infection. (Fig 3C). The C7214T mutation was also not observed in proviruses sequenced from the 9-GRev_IRES_Nef virus cultures at any of the time points examined (Fig 3A and 3C).

**Fig 3.**
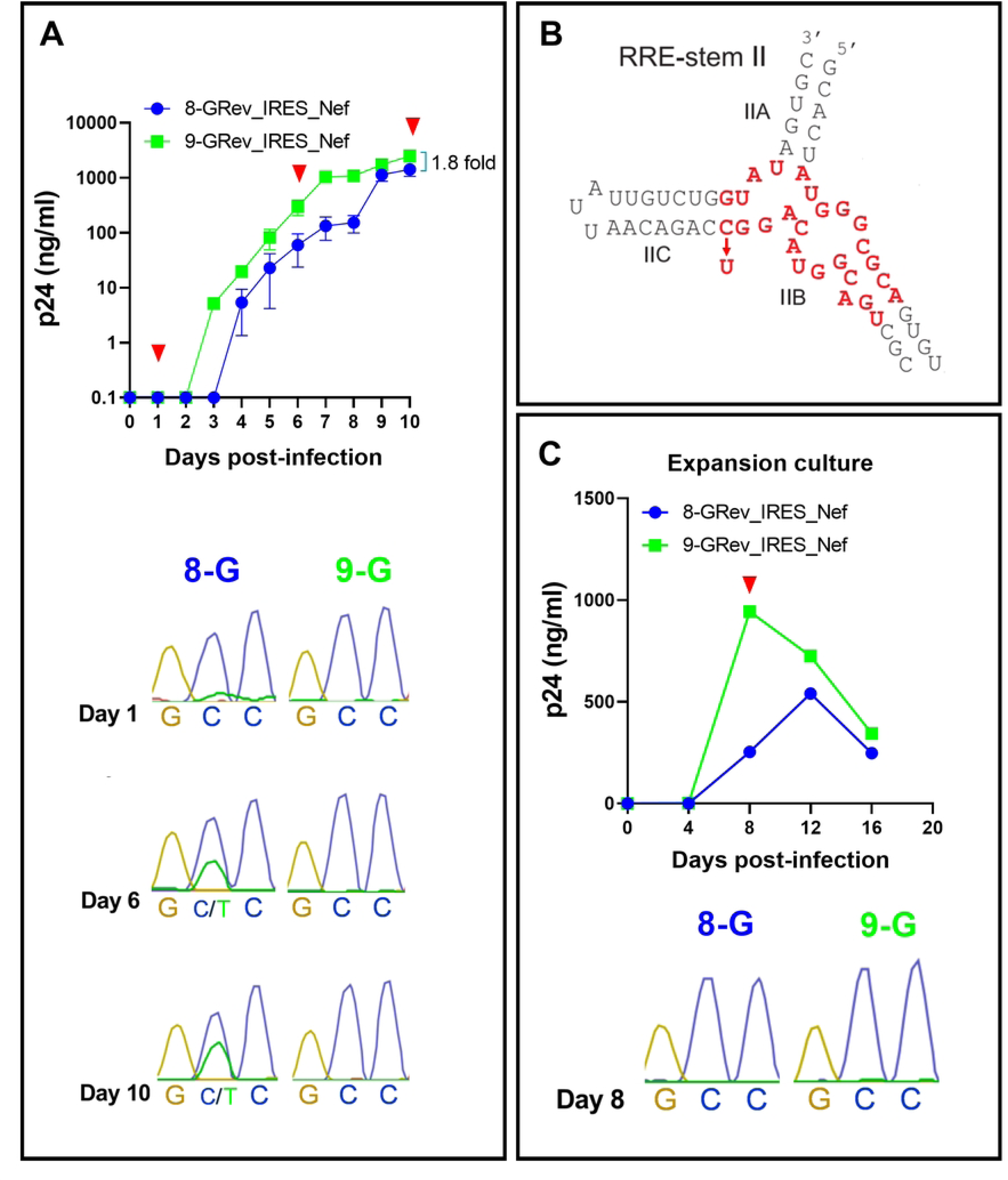
Extended passage of the low Rev activity 8-GRev_IRES_Nef virus selects for a single nucleotide change in the RRE that enhances viral replication. (A) Extended replication kinetics of 8-GRev_IRES_Nef and 9-GRev_IRES_Nef viruses in SupT1 cells. Top: Replication curve showing p24 levels measured by ELISA over time. Red arrows indicate timepoints when the RRE was sequenced from the culture cells. Bottom: Sanger sequencing chromatograms of the RRE region from infected cells at days 1, 6, and 10 post-infection, revealing the emergence of the C7214T mutation. (B) Schematic representation of the HIV-1 RRE stem II region, highlighting the location of the C7214T mutation observed in the 8-GRev_IRES_Nef virus (structure adapted from(34)). The binding site of Rev is highlighted in red, and an arrow shows the position of the C7214T mutation. (C) Characterization of the viral expansion stock. Top: Growth curve of the viral expansion stock in SupT1 cells, showing p24 levels measured by ELISA for 8-GRev_IRES_Nef and 9-GRev_IRES_Nef viruses. Bottom: Sanger sequencing chromatograms of the RRE region from SupT1 bulk culture cells used to produce viral expansion stocks at day 8 post-infection. Error bars in growth curves represent mean ± SEM of three independent experiments. Sequencing chromatograms are representative of three independent experiments.

These results suggest that the C7214T change in the RRE of the 8-GRev_RES_Nef virus, which weakened base pairing in an important region of the RRE, was selected during viral passaging because it promoted Rev-RRE function and thus enhanced replication.

### The C7214T mutation in the RRE enhances Rev-RRE dependent viral gene expression and viral replication

Since the RRE overlaps the coding region of Env, the C7214T mutation also introduced a nonsynonymous change of an alanine to a valine in gp41 (A539V). It was therefore of interest to verify that the C7214T mutation resulted in enhanced RRE function independent of a potential effect of the change in Env.

To investigate this, a fluorescence-based assay of Rev-RRE function that we have previously described was used (35). Briefly, the assay system utilizes an HIV reporter construct derived from NL4-3. This reporter contains an RRE, but does not express a functional Rev. It also contains a sequence encoding GFP inserted into the *gag* region and a sequence encoding mCherry inserted into the *nef* region. Since the GFP sequence is only present in unspliced mRNA, its expression is completely dependent on Rev-RRE activity. However, since mCherry is made from a completely spliced mRNA, its expression is Rev-RRE-independent. Additionally, the RRE in this reporter is flanked by restriction enzyme sites, allowing for easy replacement. Rev can be delivered in *trans* and Rev-RRE activity is then quantified by determining the mean fluorescence intensity of GFP in mCherry positive cells using flow cytometry.

Reporter constructs containing either the native NL4-3 RRE or the C7214T mutant RRE were constructed, along with plasmids expressing 8-G or 9-G Rev variants. In parallel experiments, the reporter plasmids were then transfected into 293T cells together with varying amounts of the 8-G Rev (Fig 4A) or 9-G Rev plasmids (Fig 4B), and the mean fluorescence intensity (MFI) of GFP was determined. For both Rev proteins, linear dose-response curves were observed. Significantly greater activity was seen with the C7214T mutant RRE compared to the original NL4-3 RRE in conjunction with both 8-G and 9-G Rev proteins, but the increase was more pronounced when 8-G Rev was expressed. This confirmed that the RRE with the C7214T mutation showed higher Rev-RRE activity, particularly in conjunction with 8-G Rev. This strengthens the notion that the mutant was selected because of higher Rev-RRE activity in the resulting 8G-Rev virus.

**Fig 4.**
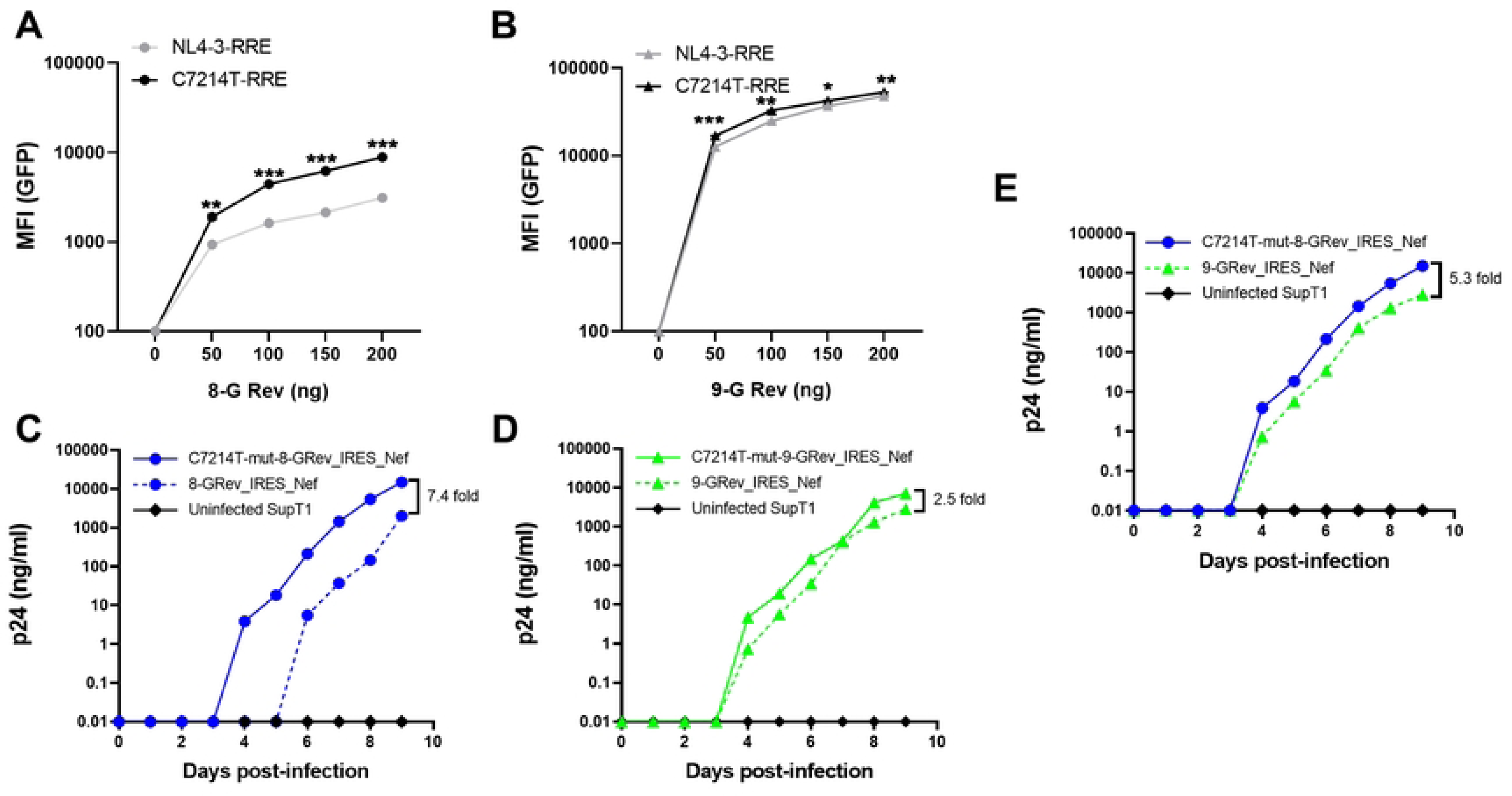
The C7214T mutation in the RRE enhances Rev-RRE dependent viral gene expression and viral replication. (A) 293T cells were co-transfected with a GFP reporter construct containing either the wild-type NL4-3 RRE or the C7214T mutant RRE, along with varying amounts of 8-G Rev expression plasmid. GFP mean fluorescence intensity (MFI) was measured by flow cytometry. (B) The Rev-RRE functional assay was performed as in (A) using 9-G Rev. (C) SupT1 cells were infected with 8-GRev_IRES_Nef and C7214T-mut8-GRev_IRES_Nef viruses at equal MOIs. Viral replication was monitored by p24 ELISA of culture supernatants over time. (D) Replication kinetics of 9-GRev_IRES_Nef and C7214T-mut9-GRev_IRES_Nef viruses were performed as in (C). (E) Comparison of replication kinetics between C7214T-mut8-GRev and 9-GRev_IRES_Nef viruses. SupT1 cells were infected with equal MOIs of each virus. Viral replication was monitored by p24 ELISA of culture supernatants over time. Error bars in panels A and B represent mean ± SD from three independent experiments. Statistical analysis was performed using unpaired, two-tailed t-test with FDR correction for multiple comparisons (**p<0.01, ***p<0.001, ****p<0.0001).

To directly assess the effect of the RRE mutation on replication kinetics, two new proviral constructs were created, introducing the C7214T mutation into both the RRE of 8-GRev_IRES_Nef (HamRek archive #6468), and 9-GRev_IRES_Nef (HamRek archive #6470) constructs. The plasmids were transfected into 293T cells to produce viral stocks, and the resulting viruses, together with the original 8-GRev_IRES_Nef, and 9-GRev_IRES_Nef viruses, were used to infect SupT1 cells. The results showed that introducing the C7214T mutation in the RRE of the 8-GRev_IRES_Nef virus greatly enhanced replication (Fig 4C), so that it now replicated better than the 9-GRev_IRES_Nef virus (Fig 4E). In contrast, this mutation had only a modest effect on the replication of the 9-GRev_IRES_Nef virus (Fig 4D). These results provide further evidence for the hypothesis that the single RRE nucleotide change was selected since it allowed increased replication of a virus expressing a Rev protein with low activity.

### High Rev-RRE activity viruses exhibit enhanced reactivation from latency and viral release compared to low Rev-RRE activity viruses

Finally, we investigated the effect of different levels of Rev-RRE activity on viral reactivation from latency using a model previously described by Lewin, with some modifications (see Methods) (Fig 5A). Resting primary CD4+ T cells were infected in parallel with either the 9-GRev_IRES_Nef, 8-GRev_IRES_Nef, or NLRev_IRES_Nef viruses, at an equal MOI. This was verified by measuring the amount of integrated provirus found in cells 24 h after infection (Fig 5B). On day three post-infection, cells were divided into two groups that were either treated with the combination of phytohaemagglutinin (PHA) and interleukin 2 (IL-2) latency reversing agents (LRA) or left untreated. Culture supernatants were collected through day 17 post-infection and analyzed for p24 production, as measured by ELISA (Fig 5A).

**Fig 5.**
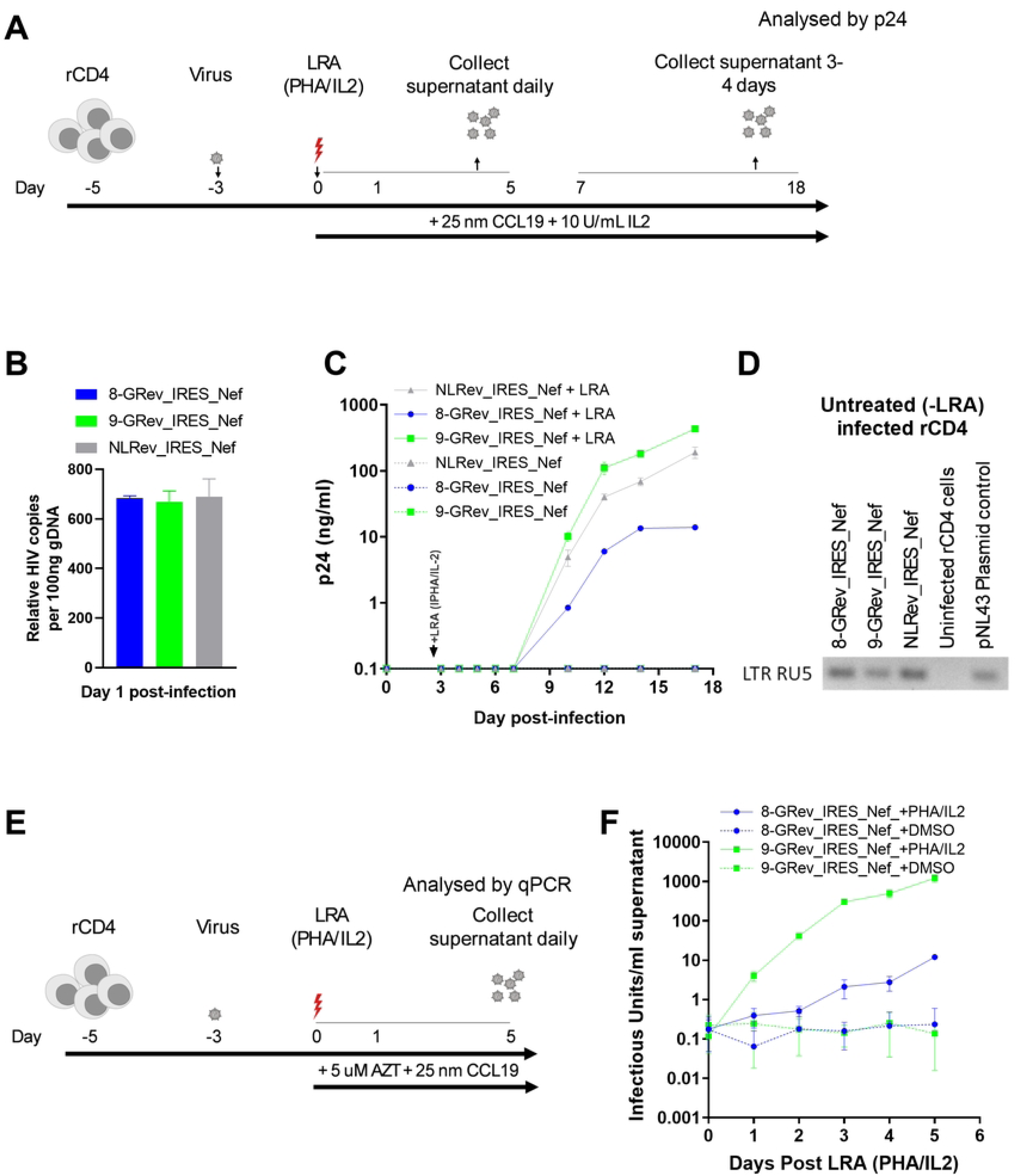
High Rev-RRE activity viruses exhibit enhanced reactivation from latency and increased viral release compared to low Rev-RRE activity viruses. (A) Schematic of the latency model and reactivation experiment. Resting primary CD4+ T cells were infected with 8-GRev_IRES_Nef, 9-GRev_IRES_Nef, or NLRev_IRES_Nef viruses. On day 3 post-infection, cells were treated with latency reversing agents (LRA: PHA + IL-2) or left untreated. Culture supernatants were collected through day 18 post-infection. (B) Quantification of integrated provirus in infected resting CD4+ T cells 24 hours post-infection, measured by Alu-gag PCR. (C) Viral reactivation kinetics following LRA treatment. P24 levels in culture supernatants were measured by ELISA over time. (D) Confirmation of integrated provirus in non-reactivated cells. Semi-nested Alu-gag PCR followed by LTR-RU5 amplification was performed on genomic DNA from cells not treated with LRA at day 18 post-infection. (E) Schematic of the reactivation experiment without reinfection. Resting CD4+ T cells were infected with 8-GRev_IRES_Nef or 9-GRev_IRES_Nef viruses, treated with LRA or vehicle at day 3, and AZT was added to prevent reinfection. (F) Viral RNA levels in culture supernatants following LRA treatment, measured by qRT-PCR. Data represents mean ± SD from three independent experiments for panel B, mean ± SD from two independent experiments for panel C, and panel F.

Following LRA treatment, cells infected with the high Rev-RRE activity virus, 9-GRev_IRES_Nef, produced p24 more rapidly and at a higher rate than cells infected with the low Rev-RRE activity 8-GRev_IRES_Nef virus (Fig 5C). The NLRev_IRES_Nef virus exhibited an intermediate behavior. As expected, cells infected with these viruses without LRA treatment, did not produce detectable p24.

To confirm the presence of integrated virus in the cells that were not reactivated, a semi-nested Alu-gag PCR followed by LTR-RU5 amplification of the first-round PCR product was performed using genomic DNA. This confirmed that the cells were infected and contained integrated proviral DNA and that the state of latency was maintained in the absence of LRAs (Fig 5D).

To assess the effect of Rev-RRE activity differences on reactivation from latency without the compounding effects of reinfection, resting primary CD4+ T cells were again latently infected with 8-GRev_IRES_Nef or 9-GRev_IRES_Nef viruses. Three days post-infection, cells were treated with LRAs or DMSO, and zidovudine (AZT), a nucleoside reverse transcriptase inhibitor, was added to prevent reinfection (Fig 5E). Supernatants and cells were collected through day 5 post-activation for analysis.

Following LRA treatment, it was observed that cells infected with the high Rev-RRE activity virus (9-GRev_IRES_Nef) released significantly more virus into the supernatant compared to the low Rev-RRE activity virus (8-GRev_IRES_Nef) (Fig 5F). Although the 8-GRev_IRES_Nef virus-infected cells exhibited some viral production post-activation, the levels were low and release was slow. Infected cells treated with DMSO alone did not reactivate as shown by only a background qRT-PCR signal.

To confirm that the difference in viral RNA production was not due to differences in cell viability, cell viability was measured in all conditions using trypan blue staining from day 1 to 5 post-reactivation (S2 Fig S2). This analysis showed no significant difference in cell viability between 8-GRev_IRES_Nef and 9-GRev_IRES_Nef infected cells (S2 Fig). Taken together, these experiments demonstrated that high Rev/RRE activity promoted latency reversal and virus production from latently infected cells.

## Discussion

In this study, we show that variations in Rev-RRE functional activity, previously observed among primary HIV-1 isolates (23), have significant impacts on both viral replication kinetics and reactivation from latency. Using engineered viruses with different Rev-RRE combinations, we found that higher Rev-RRE activity confers an advantage in viral replication and also leads to more efficient reactivation from latency in primary CD4+ T cells.

In previous work, we showed significant variation in Rev-RRE activity using Rev proteins and RREs from primary HIV-1 isolates in reporter assay(23). The experiments presented here show that activity variation also has significant effects on viral replication. Using reconstructed proviral clones, we showed significant differences in replication between viruses with different Rev and RRE sequences and that a high Rev-RRE activity virus outcompetes a low activity virus in co-infections. In addition, a spontaneous compensatory mutation in the RRE of the low Rev activity virus (8-GRev_IRES_Nef) was selected during passaging. This single nucleotide change in the RRE, within the initial Rev binding region, was sufficient to rescue the replication defect of the low-activity Rev virus and make its replication kinetics comparable to the high-activity Rev virus. This indicates that viruses with high Rev-RRE activity have a replication advantage when HIV is passaged *in vitro*. This is in stark contrast to the *in vivo* situation, where previous studies indicate that viruses with many varying levels of Rev-RRE activity are selected.

In previous work, we also analyzed Rev-RRE evolution *in vivo* (15). In individuals living with HIV, but not on antiretroviral therapy, we observed a trend towards higher Rev-RRE activity in circulating viral variants over time, driven by changes in the RRE. In one individual, the accumulation of five nucleotide changes in the RRE over several years correlated with altered RRE secondary structure as well as increased functional activity (16). The current study provides further evidence for the rapid selection of such mutations under conditions where Rev activity is limiting, emphasizing the biological significance of small RRE sequence changes.

Since the RRE also overlaps the envelope protein coding region, the C7214T mutation also led to an alanine to valine change in gp41, immediately adjacent to the start of the HR1 region. This mutation created an RRE with increased activity in our reporter assay, especially when tested with the 8-G Rev protein. This finding, combined with the fact that the mutation was only selected in the 8-GRev virus, strongly suggests that it was selected because it enhanced Rev-RRE function and not because it had an effect on gp41 activity.

Despite differences in replication kinetics, we observed that Nef expression levels were maintained at similar levels in both high and low Rev activity viruses. Nef enhances virion infectivity and is critical to maintaining high-titer replication *in vivo* (36). Nef is a potent modulator of multiple cell surface molecules, including MHC-I, CD4, CD28, CCR5, SERINC3, and SERINC5 (32, 33, 37–40). Nef also binds p53, suppressing its pro-apoptotic activity and potentially contributing to the survival of infected cells. By modulating the ratio of Nef expression to the expression of antigenic structural proteins like Gag and Env, viruses with low Rev activity are relatively protected from cytotoxic T cell-mediated killing *in vitro* (28). A specific fitness landscape could thus lead to *in vivo* selection of optimal Rev-RRE activities to achieve a balance between immune evasion, cell survival, and replication capacity.

Our experiments also showed that Rev-RRE activity levels played a significant role in viral reactivation from latency. Cells infected with the high Rev-RRE activity virus produced virus more rapidly, and at higher levels, following activation with a latency-reversing agent, compared to cells infected with the low Rev-RRE activity virus. This was observed under conditions where activation led to a spreading infection, as well as when re-infection was blocked with a reverse transcriptase inhibitor, resulting in only first-round virus production.

HIV latent virus remains the primary barrier to a cure (41) and post-transcriptional regulation of gene expression may be an important factor in latency maintenance and reversal (42). Older models of HIV persistence based primarily on transcriptional silencing have proven inadequate, as latency reversing agents designed to overcome transcriptional blocks alone are unsuccessful in fully reversing latency (43–47). Our observation that differences in Rev-RRE functional activity alter the dynamics of reactivation adds a new layer of complexity to our understanding of persistence and may serve to advance cure strategies. Our studies suggest that effective latency reversal strategies may need to consider not only transcriptional activation, but also post-transcriptional regulation mediated by the Rev-RRE axis.

The ability to significantly alter viral replication kinetics through minimal sequence changes in either Rev or the RRE provides HIV with a flexible mechanism to optimize fitness in various environments. This plasticity may allow the virus to fine-tune replication in response to selective pressures encountered during infection, such as those encountered during transmission, in different body compartments, over the course of treated or untreated HIV infection, and in entry into and emergence from latency.

## Conclusion

The results of our studies provide new insights into the complex relationship between Rev-RRE sequence variation, functional activity, and viral fitness. By demonstrating the impact of Rev-RRE activity on both replication kinetics and latency reactivation, we contribute to further understanding of how this regulatory axis affects HIV pathogenesis and persistence. These findings demonstrate the importance of post-transcriptional RNA regulation, both for HIV replication and latency, and demonstrate the importance of Rev-RRE functional variation in these processes.

## Materials and Methods

## 1. Cell Culture and Primary Cell Isolation

### 1.1 Cell Lines

HEK 293T/17 cells were cultured in Iscove’s Modified Dulbecco’s Medium (IMDM; Gibco) supplemented with 10% bovine calf serum (BCS; Cytiva) and 0.1% gentamicin (Gibco). SupT1 cells were maintained in RPMI 1640 medium with GlutaMAX (Gibco) supplemented with 10% fetal bovine serum (FBS; Cytiva) and 0.1% gentamicin. All cell lines were cultured at 37°C in a humidified incubator with 5% CO2.

### 1.2 Primary Cell Isolation and Culture

Resting CD4+ T cells were isolated from healthy donor buffy coats (American Red Cross) using the RosetteSep human CD4+ T cell enrichment cocktail (STEMCELL Technologies) according to manufacturer’s protocol. Isolated cells were maintained in RPMI 1640 medium with GlutaMAX supplemented with 10% FBS, 0.1% gentamicin, and 10 IU/ml of human interleukin-2 (hIL-2) (Boehringer Mannheim) at 37°C with 5% CO2. For latency experiments, cells were cultured with 25 nM CCL19 (R&D Systems) for 2-3 days prior to infection.

## 2. Molecular Biology and Plasmid Construction

### 2.1 HIV-1 Molecular Clones

The HIV-1 molecular clones used in this study were derived from the pNL4-3 plasmid (48). To facilitate manipulation of the Rev gene, which normally overlaps with both Env and Tat coding regions, we relocated Rev to an independent expression cassette. First, the native rev gene in pNL4-3 was functionally silenced through two modifications: the start codon was mutated from AUG to ACG, and a stop codon (UAA) was introduced at position 23 by mutating the original UAU codon. While this modification resulted in a serine to threonine substitution at amino acid position 70 in the overlapping Tat reading frame, functional assays confirmed that Tat activity was not impaired.

To enable Rev expression from an alternative location, we constructed an expression cassette containing a cDNA copy of the NL4-3 Rev gene followed by an internal ribosomal entry site (IRES). This cassette was inserted upstream of the Nef coding region, allowing both Rev and Nef to be expressed from what typically serves as the fully spliced mRNA encoding only Nef. To facilitate future modifications, the Rev sequence was flanked by unique restriction enzyme sites (*BamHI* downstream of Env and *MluI* upstream of the IRES).

Using this modified backbone, we generated two additional constructs incorporating Rev variants with different functional activities. The Rev sequences were obtained from previously characterized HIV-1 isolates: variant 8-G (GenBank accession FJ389367) and variant 9-G (GenBank accession JX140676), which exhibit approximately 4-5 fold difference in Rev activity when tested with the NL4-3 RRE (35). These Rev variants were cloned into the modified backbone using the engineered restriction sites. The resulting proviral clones were designated pNLRev_IRES_Nef (HamRek archive #5833), p8-GRev_IRES_Nef (HamRek archive #5830), and p9-GRev_IRES_Nef (HamRek archive #5831).

### 2.2 Fluorescence-Based Reporter System

The fluorescence-based Rev-RRE functional assay system has been previously described in detail (23, 35). The system consists of two packageable viral constructs. The Rev expression vector was derived from pMSCV-IRES-Blue FP (Addgene plasmid #52115, gift from Dario Vignali), where Rev variants (8-G, 9-G, or NL4-3) were inserted upstream of the IRES and eBFP2 fluorescent marker. The RRE-containing reporter construct (HamRek archive #HR5604) contains a CHYSEL (2A-like peptide)-GFP (eGFP) cassette in the gag reading frame, an mCherry gene in place of Nef, and multiple modifications to ensure replication incompetence, including deletion of the gag myristoylation site, deletion of the gag-pol frameshift, two mutations in the first exon of Rev, and a frameshift mutation in Env.

### 2.3 C7214T RRE Mutation Construction

An RRE sequence containing a C7214T mutation, flanked by *BsaI* and *AleI* restriction sites, was synthesized by IDT. This modified RRE sequence was cloned into two existing vectors: 8-GRev_IRES_Nef (HamRek archive #5830) and 9-GRev_IRES_Nef (HamRek archive #5831), generating C7214T-mut8-GRev_IRES_Nef (HamRek archive #6468) and C7214T-mut9-GRev_IRES_Nef (HamRek archive #6470), respectively. The C7214T mutation was also introduced into the fluorescence-based reporter construct (HamRek archive #5604). The RRE region in this construct is flanked by *XmaI* and *XbaI* restriction sites. This facilitated the introduction of the modified sequence containing the C7214T mutation. This created the plasmid pHR6144 which was verified by DNA sequencing.

### 2.4 Bacterial Propagation and DNA Preparation

All molecular clones were propagated in NEB Stable Competent *E. coli* at 30°C in LB media supplemented with 50 µg/mL *Ampicillin*. Plasmid DNA was isolated using the ZymoPURE II Plasmid Maxiprep Kit (NEB), quantified by spectrophotometry (260/280 >1.8), and verified by both restriction digest analysis and DNA sequencing across all modified regions.

## 3. Virus Production and Characterization

### 3.1 Initial Virus Production and Expansion

Initial viral stocks were generated by transfecting 293T cells with pNLRev_IRES_Nef (HamRek #5833), p8-GRev_IRES_Nef (HamRek #5830), or p9-GRev_IRES_Nef (HamRek #5831) plasmids. After 48 hours, supernatants were collected and p24 levels were quantified by ELISA. For viral expansion, SupT1 cells (5 × 10^6^ cells/ml) were infected with 100 ng p24 from the transfection supernatants in serum-free RPMI. DEAE-dextran was added to a final concentration of 8 µg/mL, and infection was facilitated by centrifugation at 380 RCF for one hour at 25°C. Following centrifugation, cells were washed with PBS and resuspended in RPMI supplemented with 10% FBS and gentamicin in 25 cm^2^ flasks. Cultures were maintained by removing half of the cells and medium every 2-3 days, with the culture volume replenished with fresh medium.

Large-scale viral stocks were generated by infecting 15 × 10^6^ SupT1 cells with 300 ng p24 from the peak viral expansion supernatant using the same infection protocol. Culture supernatants were collected every 4 days through day 16 post-infection, cleared of cells by centrifugation, and stored at -80°C. Viral replication was monitored by p24 ELISA, and stocks were titered by TCID50 assay. The day 8 stock, showing highest titer, was selected and used for subsequent experiments at an MOI of 0.005 for SupT1 cells and 0.05 for primary resting CD4+ T cells.

### 3.2 Rev-RRE Functional Analysis

Rev-RRE functional activity was assessed through co-transfection experiments in 293T cells. Cells were seeded at 2 × 10^5^ cells per well in 12-well plates in IMDM with 10% BCS. After 24 hours, media was replaced with serum-free IMDM, and cells were co-transfected using PEI with 1 µg of RRE-containing GFP reporter construct (either wild-type NL4-3 RRE or C7214T mutant RRE) and varying amounts of Rev expression plasmid (0, 50, 100, 150, or 200 ng of 8-G or 9-G Rev variants). Six hours post-transfection, media was replaced with IMDM containing 10% BCS. Cells were incubated at 37°C in 5% CO_2_ for 48 hours before flow cytometry analysis.

## 4. Analytical Methods

### 4.1 Flow Cytometry

#### CD4 Downmodulation Assay

Following infection of SupT1 cells with viral constructs (8-GRev_IRES_Nef, 9-GRev_IRES_Nef, or NLRev_IRES_Nef) at an MOI of 0.005 for 8 days, cells were harvested and washed with 1X DPBS. Cells were blocked with human serablock (Bio-Rad) and stained with Mouse-anti hCD4-PE conjugated antibody (R&D Systems) for 30 minutes at 4°C. After two washes, cells were fixed with 4% formaldehyde-PBS for 10 minutes and permeabilized with 0.5% triton X-100 for 10 minutes. Intracellular p24 was detected using anti-HIV-p24-FITC conjugated antibody (NIH HIV Reagent Program) for 1 hour at 4°C protected from light. Cells infected with a Nef-negative NL4-3 virus derived from the plasmid HamRek #1272 served as a control.

#### Rev-RRE Functional Activity Analysis

At 48 hours post-transfection, 293T cells were trypsinized and washed twice with cold PBS. Cells were analyzed using a sequential gating strategy. First forward and side scatter was used to identify viable cells. This was followed by single-cell discrimination, and selection of the mCherry and BFP double-positive population. To assess Rev-RRE functional activity, GFP mean fluorescence intensity (MFI) was determined in this double-positive population using FlowJo software.

All flow cytometry was performed using an Attune NxT with autosampler (Thermo Fischer Scientific). Single-color controls were used for compensation calculations in FlowJo software (v10.6.1, BD Biosciences).

### 4.2 PCR and Sequencing DNA Extraction

Cells were washed twice with cold PBS, and genomic DNA was extracted using the DNeasy Blood & Tissue Kit (Qiagen) following manufacturer’s instructions.

#### Alu-gag PCR for Proviral Detection

Integrated provirus was detected using a nested PCR approach. First-round PCR used 150 ng genomic DNA with forward Alu primer (5’-GCC TCC CAA AGT GCT GGG ATT ACA G-3’) and reverse HIV Gag primer (5’-GTT CCT GCT ATG TCA CTT CC-3’). PCR conditions were initial denaturation at 95°C for 2 minutes; followed by 25 cycles of 95°C for 15 seconds, 50°C for 30 seconds, and 72°C for 4 minutes 30 seconds. Second-round PCR used 2 µL of first-round product with forward LTR-R primer (5’-TTA AGC CTC AAT AAA GCT TGC C-3’) and reverse U5 primer (5’-GTT CGG GCG CCA CTG CTA GA-3’). Second-round conditions were initial denaturation at 95°C for 2 minutes, followed by 35 cycles of 95°C for 15 seconds, 50°C for 30 seconds, and 72°C for 30 seconds. PCR products were visualized by electrophoresis for the ∼100 bp RU5 product.

#### Competition Assay

SupT1 cells were co-infected in triplicate with 8-GRev_IRES_Nef and 9-GRev_IRES_Nef viruses at an MOI of 0.005 each. Cellular DNA was collected on days 1, 3, 5, and 7 post-infection. The relative proportion of integrated viruses was determined by discriminative PCR using 200 ng genomic DNA template in 50 µL reactions containing Taq DNA polymerase (Thermo Scientific), 2 pmol env forward primer (5’-CCTAGAAGAATAAGACAGGGC-3’), and 1 pmol each of variant-specific reverse primers: 8-G-specific (5’-CCCCAGATATTTCAGGCCCTC-3’) and 9-G-specific (5’-GTCTCTCAAGCGGTGGTAGCAC-3’). PCR cycling conditions were initial denaturation at 95°C for 3 minutes; 25 cycles of 94°C for 30 seconds, 60°C for 30 seconds, and 75°C for 45 seconds; followed by final extension at 72°C for 10 minutes.

PCR products (15 µL) were separated on 2% ultrapure agarose gels (Invitrogen) containing ethidium bromide at 80V for 2 hours. Gels were imaged using a Bio-Rad transilluminator with optimized exposure settings to prevent band saturation. Band intensities were quantified using Image Lab software (Bio-Rad). Standard curves were generated using known ratios of 8-GRev and 9-GRev plasmids, which were also amplified separately or together as PCR controls. The relative proportion of each variant was expressed as a percentage of total signal from both PCR bands per lane.

#### RRE and Rev Sequencing Analysis

To analyze potential mutations, both RRE and Rev regions of proviruses were amplified using nested PCR. First-round PCR reactions contained 150 ng genomic DNA template in 25 µL reactions containing: 1X PCR buffer, 1.8 mM MgSO4, 0.2 mM each dNTP, 0.2 µM each primer, and 1 U high-fidelity Taq DNA Polymerase (Invitrogen). For RRE amplification, first-round primers were 5’-GTGACACAATCACACTCCCATGC-3’ (forward) and 5’-GGTGAATATCCCTGCCTAACTC-3’ (reverse), followed by second-round primers 5’-CAATGGGTCCGAGATCTTCAGACC-3’ (forward) and 5’-CACTCCATCCAGGTCATGTTATTC-3’ (reverse). Rev amplification used first-round primers 5’-GCACTTATCTGGGACGATCTGCG-3’ (forward) and 5’-GCTTCCTTCACGACATTCAACAG-3’ (reverse), followed by second-round primers 5’-CCTACAGTATTGGAGTCAGGAAC-3’ (forward) and 5’-GGAATGCTCGTCAAGAAGACAGGGCC-3’ (reverse).

PCR conditions for both rounds consisted of initial denaturation at 94°C for 3 minutes, followed by 20 cycles of denaturation at 94°C for 30 seconds, annealing at 52°C (first round) or 53°C (second round) for 30 seconds, and extension at 68°C for 50 seconds, with a final extension at 68°C for 10 minutes. Second-round PCR used 5 µL of first-round product as template. PCR products were visualized on 2% agarose gels containing ethidium bromide, extracted and subjected to Sanger sequencing. All reactions were performed in duplicate to ensure reproducibility.

### 4.3 Protein Analysis

#### Western Blot Analysis

Infected cells were harvested and washed twice with 1X Dulbecco’s Phosphate-Buffered Saline (DPBS; Invitrogen). Cell pellets were lysed in buffer containing 1 mM sodium chloride, 1X protease inhibitor cocktail, 1 mM EDTA (pH 7.4), and 1% SDS by heating at 95°C for 10 minutes. Lysates were cleared by centrifugation (14,000 × g, 10 minutes, room temperature), and supernatants were mixed 1:1 with LDS Sample Buffer (Invitrogen) and heated at 72°C for 10 minutes. Equal volumes of protein lysate were separated on NuPAGE 4-12% Bis-Tris (MES) gels (Invitrogen) under reducing conditions and transferred to PVDF membranes using the Trans-Blot® Turbo™ Transfer System (Bio-Rad) at 30V for 1 hour.

Membranes were blocked with iBind™ Flex Solution (Thermo Fisher Scientific) for 10 minutes at room temperature. Primary antibodies (mouse anti-p24 and mouse anti-Nef monoclonal antibodies (NIH HIV Reagent Program, Division of AIDS, NIAID, NIH), both at 1:1000 dilution and secondary antibody (IRDye 800CW Goat anti-Mouse IgG, LI-COR Biosciences, 1:6000 dilution) were applied using an iBind Flex Western System (Thermo Fisher Scientific) for 2.5 hours at room temperature. Beta-tubulin served as loading control for normalization. Protein bands were visualized using an Odyssey CLx chemifluorescence detector (LI-COR Biosciences) and quantified using ImageStudio software version 5.2 (LI-COR Biosciences). All western blot analyses were performed with at least three biological replicates.

## 5. Latency Model Experiments

### 5.1 Establishment of Latent Infection

Latent infection was established using a modified version of the Lewin model. Following CCL19 treatment (described in section 1.2), latent infection was initiated by spinoculation of 2 × 10^6^ cells with HIV-1 Rev-variant molecular clones at an MOI of 0.05 for 2 hours at 300 × g at room temperature. Following spinoculation, cells were washed three times with room temperature 1X DPBS to remove residual virus and maintained in culture media supplemented with 25 nM CCL19 and 10 IU/ml IL-2.

### 5.2 Latency Reversal Analysis

Three days post-infection, latently infected cells were divided for two parallel reactivation experiments. In the first approach, cells were reactivated using 10 µg/mL phytohemagglutinin (PHA) (Sigma-Aldrich) and 10 IU/mL IL-2. Virus outgrowth was monitored for up to 18 days by p24 ELISA of culture supernatants. In the second approach, reactivation was performed in the presence of 5 nM zidovudine (AZT) (NIH HIV Reagent Program) to prevent reinfection, and virus production was monitored for 5 days post-reactivation.

### 5.3 Viral RNA Analysis and Quantification

Supernatants from AZT-treated cultures were collected daily and processed for RNA analysis. Samples were cleared of cellular debris by centrifugation (5300 RCF, 5 minutes, 4°C), and viruses were pelleted by centrifugation at 16,000 RCF for 1 hour at 4°C. Virus pellets were resuspended in 50 µL RNase-free 5 mM Tris-HCL (pH 8.0) and treated with 10 µL proteinase K (20 mg/mL) at 55°C for 30 minutes. RNA was isolated by adding 200 µL of 6 M guanidinium isothiocyanate and 10 µL glycogen (20 mg/mL), followed by isopropanol precipitation. RNA pellets were washed with 70% ethanol and resuspended in 40 µL RNase-free Tris-HCL. cDNA synthesis was performed using the SuperScript IV Reverse Transcriptase kit with oligo(dT)20 primer.

Viral RNA was quantified by qPCR using 2 µL cDNA template with the Luna TaqMan Master Mix (NEB) on an ABI Step One Plus thermocycler. Reactions contained 0.3 nM each primer (LTR-R forward: 5’-TTA AGC CTC AAT AAA GCT TGC C-3’; U5 reverse: 5’-GTT CGG GCGCCA CTG CTA GA-3’) and 0.2 nM of double quenched RU5-specific probe (5’-/56-FAM/CCAGAGTCA/ZEN/CACAACAGACGGGCACA/3IABkFQ/-3’). PCR conditions were initial denaturation at 95°C for 1 minute, followed by 40 cycles of 95°C for 15 seconds and 60°C for 30 seconds. Standard curves were generated using 10-fold serial dilutions of predetermined NL4-3 cDNA from TCID50-quantified virus, with values converted to infectious units per mL supernatant. Results were analyzed using StepOne Software v2.3. To ensure RNA specificity and exclude DNA contamination, control reactions were performed on supernatant samples without reverse transcriptase treatment. These controls confirmed the absence of contaminating HIV DNA in the supernatant nucleic acid preparations.

### 5.4 Cell Viability

Cell viability was assessed using the Trypan Blue exclusion method. Cells were mixed with an equal volume of 0.4% Trypan Blue solution (Gibco) and counted using a hemocytometer.

## 6. Statistical Analysis

Statistical analyses were performed using GraphPad Prism software version 8.0.1 (GraphPad Software, USA). For experiments with three independent replicates, statistical comparisons were conducted using unpaired, two-tailed t-tests, with false discovery rate (FDR) correction applied for multiple comparisons. P-values < 0.05 were considered statistically significant. For experiments with two independent replicates, only descriptive statistics were calculated. All data are presented as mean ± standard deviation (SD) or standard error of the mean (SEM) as indicated in figure legends.

## Acknowledgments

The flow cytometry data for this manuscript were generated at the University of Virginia Flow Cytometry Core Facility (RRid:SCR_017829), with partial support from the NCI Grant (P30-CA044579). Monoclonal antibodies were obtained through the NIH HIV Reagent Program, Division of AIDS, NIAID, NIH. The anti-HIV-1 Nef antibody (AE6, ARP-709) was contributed by Dr. James Hoxie and the monoclonal anti-HIV-1 p24 antibody (183-H12-5C, ARP-3537) was contributed by Dr. Bruce Chesebro and Kathy Wehrly. Partial salary support was provided by the Myles H. Thaler Endowed Professorship (D.R.) and the Charles Ross Jr Endowed Professorship (M.-L. H) at the University of Virginia, and NIH grant K08 AI136671 (P.E.H.J). Research support was provided by Myles H. Thaler Research Support Gift and NIH grant R21 AI134208.

